# Transcriptomic insights into triploid seed failure in *Arabidopsis arenosa* natural populations

**DOI:** 10.64898/2026.04.09.717150

**Authors:** Susnata Salony, Martin Kovačik, Vojtěch Čermák, Adéla Přibylová, Aleš Pečinka, Filip Kolář, Clément Lafon Placette

## Abstract

- Polyploidy is a key evolutionary force in plants; among its many consequences, hybridization between diploids and polyploids is restricted due to the “triploid block”. While the molecular mechanisms of this postzygotic barrier are extensively studied in the model species *Arabidopsis thaliana*, our understanding of the triploid block in natural systems remains limited.
- Here, we investigated the transcriptome of failing triploid seeds in *Arabidopsis arenosa*, a close relative of *A. thaliana* with diploid and autotetraploid populations meeting in nature. We also identified *A. arenosa* imprinted genes.
- Triploid seeds showed preferential misregulation of imprinted genes, which parallels the parent-of-origin features of the triploid block. Tissue-specific transcriptomic analyses revealed pathogen defense-like response being recurrently affected in the endosperm and seed coat. This pathway is commonly misregulated in all species with triploid seed transcriptomes studied to date. The associated genes however are likely more involved in cell-cell signaling rather than pathogen defense *per se*.
- Altogether, this study depicts a thorough molecular landscape of the triploid block for the first time in natural systems. Combining data on the understudied maternal excess transcriptome, a new imprintome, a tissue-specific focus, and a cross-species comparison, this study also advances our understanding of the triploid block and seed development.

## INTRODUCTION

Polyploidy, or whole-genome duplication (WGD), is a major driver of plant speciation and adaptation (Soltis & Soltis, 2012). It has been estimated that 15 % of speciation events in flowering plants were associated with change in ploidy (Wood et al., 2009) and when a higher extinction rates in neopolyploids are taken into account, polyploid speciation rates may be even higher up to 30% (Mayrose et al., 2011). Newly formed polyploids often become immediately reproductively isolated from their diploid progenitors, restricting gene flow and promoting genetic divergence (Ramsey & Schemske, 1998, Husband & Sabara, 2004). One of the critical mechanisms contributing to this reproductive isolation is the “triploid block”, where hybrid seeds formed between parents of different ploidy levels fail to develop properly (Scott et al., 1998; Köhler et al., 2010; Lafon-Placette & Köhler, 2016). This seed inviability is typically caused by disruptions in the strict 2:1 maternal-to-paternal genomic ratio in the endosperm, which is essential for normal seed development (Haig & Westoby, 1989). Deviations from this ratio can lead to abnormal endosperm development, such as over-proliferation or impaired cellularization, both of which ultimately result in embryo arrest (Erilova et al., 2009; Lafon-Placette & Köhler, 2016).

Much of the mechanistic understanding of the triploid block comes from studies in *A. thaliana*. In this model species, paternal-excess crosses (2x × 4x; the seed parent is always designated first) often cause delayed endosperm cellularization leading to seed collapse, whereas maternal-excess crosses (4x × 2x) typically result in precocious endosperm cellularization and mostly viable seeds (Scott et al., 1998). Because of its high viability, maternal-excess cross is usually neglected in *A. thaliana* studies except for early studies (Scott et al., 1998; Tiwari et al., 2010), leading to a significant gap in our understanding of the genetic basis of triploid block as a whole, particularly from a genome-wide perspective. In addition, high viability of maternal-excess seeds is not the rule, with many known cases showing uniformly very strong triploid block in both directions of crosses (Sekine et al., 2013; Vallejo-Marin et al., 2016; Salony et al., 2025). In fact, a substantial rate of lethality was observed in maternal-excess seeds even in the closely related outcrossing species *A. arenosa* (Morgan et al., 2021). These parent-of-origin specific defects highlight the dosage sensitivity of the endosperm being shaped by genomic imprinting—an epigenetic mechanism regulating gene expression based on parent-of-origin. In plants, genomic imprinting predominantly occurs in the endosperm, where certain genes are expressed preferentially or exclusively from either the maternal or paternal allele (Köhler et al., 2012; Gehring, 2013). The Polycomb Repressive Complex 2 (PRC2), which includes *MEA*, *FIS2*, and *FIE*, plays essential roles in regulating genomic imprinting, and its knock-out mutations phenotypically resemble the defects of paternal-excess triploid seeds (Chaudhury et al., 1997). In contrast, disrupting paternally expressed genes leads to viable paternal-excess triploid seeds in *A. thaliana* (Dilkes et al., 2008; Kradolfer et al., 2013; Wolff et al., 2015), bringing a causal link between imprinted genes and the triploid block. While molecular studies focus on the endosperm when it comes to the triploid block, other seed compartments are relatively understudied. Recently, the transcriptional response of the embryo, a direct victim of endosperm defects in triploid seeds, has been studied in paternal-excess seeds (Xu et al., 2023). Additionally, few studies have shown that altering seed coat genes can rescue paternal-excess seeds, suggesting that the seed coat may also play a role in the triploid block (Zumajo-Cardona et al., 2023). However, how the embryo and seed coat respond to the triploid block, particularly in maternal-excess crosses, remains poorly understood.

The molecular basis of the triploid block and its link with genomic imprinting is thus well-established in the thoroughly-studied species *A. thaliana*. Transcriptomic studies have been conducted in other systems such as *Brassica* and rice (Stoute et al., 2012; Wang et al., 2018), building premises of a broader mechanistic view on the triploid block. However, all these cases involve comparison of diploid model lines with artificial tetraploids derived from them. It is unclear to what extent this knowledge applies to species where the triploid block actually plays a role in nature, in other words, to species showing natural polyploid populations and contact zones between diploids and polyploids where both ploidy levels meet and reproductively interact.

*A. arenosa*, a close relative of *A. thaliana*, offers an ideal natural system to study how reproductive barriers evolve post-polyploidization. *A. arenosa* tetraploids originated approx. 20-30 thousand generations ago from their diploid ancestors in Western Carpathians (present day Slovakia), where both ploidies still co-occur, forming a primary contact zone (Arnold et al., 2015; Monnahan et al., 2019). Tetraploid populations then spread across Europe and came into contact with additional, earlier diverging diploid lineages of the same species in different regions, forming additional secondary contact zones in the Southeastern Carpathians (Romania) and northern Dinarides (Slovenia) (Morgan et al., 2020, 2021).

Interestingly, despite contacts between the cytotypes, no triploid individuals can be observed in nature in these contact zones (Morgan et al., 2021). Paternal-excess is virtually fully lethal, while maternal-excess seeds show substantial lethality rates too (Morgan et al., 2021), likely contributing to the high reproductive isolation observed between cytotypes in the wild. Interestingly, while triploid seeds display similar phenotypic defects over different contact zones, they also show natural variation in the viability, thus providing natural biological replicates to study the triploid block in the wild. In this study, we investigate hybrid seed failure between diploid and tetraploid *A. arenosa*, using populations sampled across two contact zones—the Southeast Carpathians (SEC; secondary contact) and the West Carpathians (WCA; primary contact)—to capture a broader spectrum of natural variation corresponding with varying levels of inter-ploidy divergence and different historical evolutionary context. Using reciprocal interploidy crosses (paternal – 2x × 4x, and maternal – 4x × 2x – excess crosses), we analyze the overall transcriptional signatures and specific imprinted gene expression patterns associated with hybrid seed failure. Specifically, we ask: (1) What biological pathways are affected by the triploid block in *A. arenosa*? (2) Are imprinted genes prominent players in the triploid block transcriptional response? (3) What is the transcriptomic response of each seed compartment (seed coat, embryo, endosperm) to the triploid block?

To address these questions, we first identified transcriptomic profiles of individual seed tissues—endosperm, embryo, and seed coat— and inferred imprinted genes in diploid *A. arenosa*. Then, we performed interploidy crossings in representatives from two natural ploidy contact zones of *A. arenosa*, generated whole-seed transcriptomes, characterized non-additive expression patterns in interploidy hybrid seeds as compared to homoploid controls, and identified candidate genes for triploid block. By combining tissue-specific and whole-seed transcriptomic analyses, our study aims to characterize the molecular basis of interploidy barriers in natural contact zones of *A. arenosa* and understand how these mechanisms contribute to reproductive isolation and polyploid speciation in the natural plant system.

## MATERIALS AND METHODS

### Plant material

For the identification of imprinted genes, seeds of *A. arenosa* were collected between 2018 and 2022 from 14 natural populations across its natural range in Central and Southern Europe, including representatives of the Western Carpathian, Pannonian, and Dinaric lineages. Populations belonging to divergent (but still cross-compatible) diploid lineages were selected to maximize the number of SNPs between parents and enable parent-of-origin sequence detection. The GPS locations are provided in Supp Table 3.

For the triploid block experiment, sympatric diploid and tetraploid individuals from two distinct ploidy contact zones of *A. Arenosa* were used: the Western Carpathian (WCA – Slovakia, primary contact zone, GPS: 2x – 48.96221N, 20.40265E, 4x – 48.95614N, 20.41459E, AA372 of Morgan et al., 2021, Fig. S3) where genetically close diploid and autotetraploid plants meet (PHD–PHT, Fst = 0.0449978; Bohutinska et al., 2024) and the South-Eastern Carpathian (SEC – Romania, GPS: 2x – 45.53349N, 25.29145E, 4x – 45.52998 N, 25.26694 E, AA250 of Morgan et al., 2021) representing secondary contact of genetically divergent ploidy cytotypes (BUD–BUT, Fst = 0.101037; Bohutinska et al., 2024). To generate plants for the crossing experiment, at least 20 seeds per ploidy and contact zone were sown. These seeds, randomly selected irrespective of their shape or size, originated from five seed parents, each of diploid and tetraploid, obtained in control of homoploid crosses from a previous crossing experiment with the same lineages (Morgan et al., 2021).

### Growth conditions

Prior to germination, seeds were pre-treated by incubation at 37 °C for 2 days, followed by −18 °C for 2 days to minimize the risk of thrips infestation. Seeds were surface-sterilized using a solution containing 5% sodium hypochlorite and 0.01% (v/v) Triton X-100, then plated on Petri dishes containing Murashige and Skoog medium (1/2× MS salts (DUCHEFA BIOCHEMIE B.V M0222), 0.8% plant agar (DUCHEFA BIOCHEMIE B.V P100), 10 mM MES (DUCHEFA BIOCHEMIE B.V M150), pH 5.8). Seeds were stratified for 7 days at 4 °C in the dark and then germinated under controlled conditions (21/18 °C with a 16/8 hour day/night cycle) for two weeks. Germinated seedlings were transferred to soil and grown under the same conditions for five additional weeks, followed by vernalization at 4 °C under short-day conditions (8/16 hour day/night cycle) for at least two months. After vernalization, plants were returned to long-day conditions (21/18 °C with a 16/8 hours day/night cycle) to induce flowering.

### Flow cytometry

We determined the ploidy level of all plants involved in the study using flow cytometry. Nuclei were isolated from fresh leaf tissue using Otto I buffer (0.1 M citric acid, 0.5% Tween-20; Otto and Whitton, 1990) following a two-step protocol (Doležel et al., 2007), with *Carex acutiformis* (2C = 0.82 pg; Lipnerová et al., 2013) as an internal standard. Nuclear suspensions were filtered through a 42 μm nylon mesh and stained with Otto II buffer (0.4 M Na2HPO4·12H2O) supplemented with β-mercaptoethanol and DAPI (final concentration: 4 μL mL−1). Samples were run on a Partec CyFlow ML flow cytometer (Partec GmbH, Münster, Germany, UV LED diode), with a minimum of 1500 events recorded per sample. Histograms were processed using FloMax FCS 2.0 (Partec), and only those with G0/G1 peak CVs below 5% were included in the analysis.

### Crossing designs

For the identification of imprinted genes, to maximize the number of SNPs between parents, crosses were performed between plants of different populations (in total 26 crosses—13 reciprocal pairs, between 14 diploid populations were performed, see Supp Table 4 for the details of the crossing design). A structured pairwise reciprocal crossing design was implemented, in which selected individuals from different populations were crossed in both directions (i.e., as seed parent and pollen donor) to capture parent-of-origin effects. Crosses were done by emasculating the unopened flower buds of the seed parent and transferring pollen from the pollen donor 1-3 days after emasculation to the pistil.

For the triploid block experiment (Fig. S3, Supp Table 5), a total of 40 plants were used from both WCA (2x, n = 10; 4x, n = 9) and SEC (2x, n = 10; 4x, n = 10) contact zones. Four types of cross-treatment were performed: (1) interploidy cross with diploid seed parent and tetraploid pollen donor (2x × 4x), (2) interploidy cross with tetraploid seed parent and diploid pollen donor (4x × 2x), (3) diploid control (2x × 2x), and (4) tetraploid control (4x × 4x). Treatments 1 and 2 were reciprocal, with the same individuals used as both seed parent and pollen donor. Manual pollinations were done as explained in the previous paragraph.

### Endosperm, embryo, seed coat and whole-seed collection

For the identification of imprinted genes, siliques were collected 10-16 days after pollination, and endosperm, embryo, and seed coat were manually isolated and separated in the developmental stage, varying from early to late torpedo stage, all at room temperature (∼24 °C). Siliques were placed on 3M micropore tape, opened by tweezer (Precision tweezers DUMONT® straight with extra fine tips Dumoxel®, 5) and individual seeds were transferred into extraction buffer (0.3 M D(-)-Sorbitol (Millipore® 1.07758), 5 mM MES (DUCHEFA BIOCHEMIE B.V M1503), pH 5.7, filter sterilized). Using a tweezer and insulin syringe (Omnican® 100), individual parts of the seeds were separated. Endosperm and embryo samples originated from 120-160 seeds, and seed coat samples from 240-320 seeds per cross. Individual fractions were transferred into 2.0 ml safe-lock tubes (Eppendorf 0030123344) and centrifuged for 90 sec at 12,000 g in the bench MiniSpin® centrifuge. Supernatant from all samples was discarded. Endosperm samples were frozen immediately in liquid nitrogen, and embryos and seed coats were washed two times in 1 ml of extraction buffer before freezing. All samples were kept at −80 °C for further use.

For the triploid block experiment, developing siliques were collected 14 days after pollination (DAP 14), corresponding to embryo stages 5 (heart) to 6 (torpedo), which coincides with significant phenotypic differences observed between crosses (Morgan et al., 2021). We collected 5 biological replicates per cross-type per contact zone (5 maternal-excess interploidy, 5 paternal-excess interploidy, 5 diploid control, 5 tetraploid control crosses per contact zone; total n = 40). Each replicate consisted of approximately 80–100 seeds harvested on the same seed parent and pollinated by the same pollen donor (∼10–15 siliques). Siliques were harvested into sterilized Eppendorf tubes, flash-frozen in liquid nitrogen, and stored at −80 °C until RNA extraction.

### RNA and DNA sequencing

RNA was isolated using the RNAqueous™ Total RNA Isolation Kit (Thermo Fisher Scientific AM1912) with the inclusion of Plant RNA Isolation Aid (Thermo Fisher Scientific AM9690) in the lysis buffer. DNA was extracted from leaves using the DNeasy® Plant Mini Kit (Qiagen 69104).

Total RNA and DNA samples were sent to Novogene (UK) Company Limited for library preparation and sequencing. RNA samples undergo quality control, poly A enrichment, directional mRNA library preparation, and Illumina Sequencing PE150, with raw data obtained 4.5 G and 7.5 G. DNA samples undergo quality control, whole genome library preparation (350 bp), and Illumina Sequencing PE150, with raw data obtained 6 G.

### Gene annotation

Structural gene annotation of *A. arenosa* genome assembly (GCA_026151155.1; Bramsiepe et al., 2023) was done using Augustus version 3.3.2 (Stanke et al., 2008). The full-length coding sequences from *Arabidopsis lyrata* and training annotation files of *A. thaliana* were used for homology-based gene prediction, resulting in 33,911 gene models. Functional annotation of predicted protein sequences was done using 21,867 and 23,926 reciprocal blast hits (e-val less than 0.001) from *A. thaliana* and *A. lyrata*, respectively.

### Variant calling and identification of imprinted genes

Paired-end 150 bp sequenced reads from both parental DNA and offspring RNA were mapped to the *A. arenosa* reference genome (GCA_026151155.1) using BWA version 0.7.17 (Li, 2013) with a smart pairing option for paired-end alignment. Duplicates were then removed and SNPs called using GATK version 4.2.5.0 (McKenna et al., 2010). The obtained SNPs were filtered by genome coverage (DP > 7) and genotype quality (GQ > 19). SNPs within exons and without a common parental allele were further used for the identification of imprinted genes. Each SNP was tested for significant (p-adj < 0.05) deviation from the expected ratio of maternal to paternal allele 2:1 in both reciprocal crosses using the Binomial test. Imprinted genes were identified when a gene contained at least two SNPs showing significant parent-of-origin bias in the same direction. Additionally, we took into consideration that at least two-thirds of all informative SNPs within the gene (regardless of significance) exhibited the same parental bias.

Finally, we removed genes that might represent contamination from maternal tissue rather than imprinted genes. To do so, the raw reads from endosperm and seed coat samples were trimmed for adaptors using Trim Galore version 0.4.1 (Martin, 2011) and aligned to the *A. arenosa* reference genome (GCA_026151155.1) using HiSat2 version 2.1.0 (Kim et al., 2019). Reads were then assigned to features and meta-features using Subread version 1.5.2 (Liao et al., 2013) according to the prepared genome annotation. Differential expression analysis was performed using DESeq2 version 1.38.3 (Love et al. 2014) in R v.4.2.2 (R Core Team 2020) to compare gene expression between endosperm and seed coat samples. Genes identified as parentally biased and significantly upregulated in endosperm compared to seed coat (Benjamini–Hochberg FDR-adjusted P-value < 0.05) were kept as imprinted genes.

### Differential gene expression analysis for triploid seeds

Quality control of raw reads was performed with FastQC for the whole-seed transcriptomics data. Adapters and low-quality reads were trimmed using FastX and Trimmomatic with the following parameters: SE.fa:2:30:10, ILLUMINACLIP:TruSeq3, LEADING:3, TRAILING:3, SLIDINGWINDOW:4:15, MINLEN:36. Trimmed reads were aligned to the *A. arenosa* reference genome (GCA_026151155.1) using STAR alignment with default settings and additional parameters. Uniquely mapped reads were quantified using HTSeq-count (Anders et al., 2015) with parameters: --stranded=no. The *A. arenosa* genome annotation explained above was used for feature counting. Read counts were processed using DESeq2 in R for differential expression analysis (Love et al., 2014).

Before differential expression analysis (DEA), read counts from all 40 seed samples were normalized for sequencing depth and composition using the variance-stabilizing transformation (vst) function in DESeq2 package (Love et al., 2014). Normalized sample-level expression profiles were visualized using principal component analysis (PCA) via the plotPCA function. Although the initial dataset consisted of 40 samples, one sample from the WCA population was excluded from further analyses due to outlier behavior in the PCA, resulting in a final dataset of 39 samples. Since low-count filtering did not affect trnascriptome-wide patterns (PCA), DEA was performed without prior filtering. Independent filtering based on the mean of normalized counts was applied by default in the DESeq2 results function. To account for potential lineage-specific (WCA vs SCA contact zone, batch) effects, differential expression models were fitted using a design formula that included both cross-treatment (target contrast) and contact zone (batch) as factors. A custom ‘contraster’ function (https://www.atakanekiz.com/technical/a-guide-to-designs-and-contrasts-in-DESeq2/) was used to compare each interploidy hybrid to the average expression of both diploid and tetraploid controls within the same contact zone. These averages were calculated at the population level, not paired by families, enabling identification of genes that were non-additive relative to the typical expression range of each lineage. Genes with |log2FoldChange| ≥ 1 and adjusted P < 0.05 were considered differentially expressed, with log2FoldChange values calculated relative to the average expression of the diploid and tetraploid controls within the same contact zone. This enabled us to identify genes exhibiting non-additive expression in interploidy hybrids, relative to parental expressions.

### Tissue-specific gene expression deconvolution

First, we estimated tissue composition within whole-seed RNA-seq samples by performing gene expression deconvolution using CIBERSORTx (Newman et al., 2019), based on a set of tissues-specific genes obtained from manually dissected and sequenced embryo, endosperm, and seed coat tissues. To identify tissue-specific genes, pairwise differential expression analyses between each tissue type were performed using DESeq2. Genes upregulated in all pairwise comparisons (log2FC ≥ 1, adjusted P < 0.05) were classified as either endosperm-, embryo-, or seed coat-specific. This reference tissue-specific dataset was used to generate a reference signature matrix. The signature matrix was constructed on the CIBERSORTx web platform with quantile normalization disabled to preserve raw expression distributions. This matrix was used to impute tissue fractions in each of the 39 bulk RNA-seq samples based on whole-seed expression data. Analysis was run with S-mode batch correction to minimize technical discrepancies between the reference and bulk datasets, and 100 permutations were applied to assess confidence in the estimated tissue proportions. Next, we inferred tissue-specific gene expression profiles by employing the High-Resolution mode of CIBERSORTx using its Docker implementation. Through this, we obtained tissue-level gene expression profiles for ∼30,000 genes, separately inferred for embryo, endosperm, and seed coat in every bulk RNA-seq sample. S-mode batch correction was again applied to account for technical variability between the reference and bulk datasets. The resulting tissue-specific expression matrices were further analyzed using DESeq2, following the same pipeline described above for the whole-seed transcriptome data (see above).

### GO enrichment analysis

Gene ontology (GO) enrichment was performed using the PlantRegMap web tool (Tian et al., 2019), with *A. thaliana* homologs used both for annotation and as the background gene list (Supp Dataset 10), to reduce interspecies bias and improve interpretability. Enrichment was tested using Fisher’s exact test, with Benjamini–Hochberg correction for multiple testing (FDR < 0.05). q-values (−log₁₀ transformed) were visualized in dot plots grouped by GO category using the ggplot2 package in R. Dot size reflected the number of DEGs per term, and color gradient corresponded to adjusted significance level.

### Analysis of imprinted gene expression in triploid block samples

The resulting list of 96 imprinted genes (47 MEGs, 49 PEGs) was intersected with the set of differentially expressed genes (DEGs) identified from whole-seed RNA-seq using the GeneOverlap package in R v4.3.1, which applies Fisher’s exact test to determine significant overlap between gene sets.

Raw read counts were normalized to transcripts per million (TPM), and principal component analysis (PCA) was performed on expression values of imprinted genes across all 40 samples using the ‘prcomp’ function in R.

### Accession numbers

Accession numbers for RNA and DNA data generated for both the imprinting analysis and the triploid block experiment are provided in the Data Availability statement.

## RESULTS

### Overall gene expression patterns across the cross types in the triploid block experiment

To explore the molecular landscape of triploid seed failure in *A. arenosa*, we performed a PCA on TPM-normalized gene expression profiles. PC1 (20.16%) separated samples by contact zones, forming two distinct clusters corresponding to WCA and SEC contact zones. PC2 (11.81%) showed a segregation according to the cross type (Fig. 1A). Within each contact zone cluster, interploidy hybrids (2x × 4x and 4x × 2x) formed separate sub-clusters at the extreme ends of the gradient, with their respective diploid and tetraploid controls occupying intermediate positions in between them. PERMANOVA analysis confirmed this pattern, showing that contact zone explained the largest proportion of variance (29%), followed by cross type (25%) and their interaction (64%), all statistically significant (p = 0.001) (Supp Table 1).

**Fig. 1.**
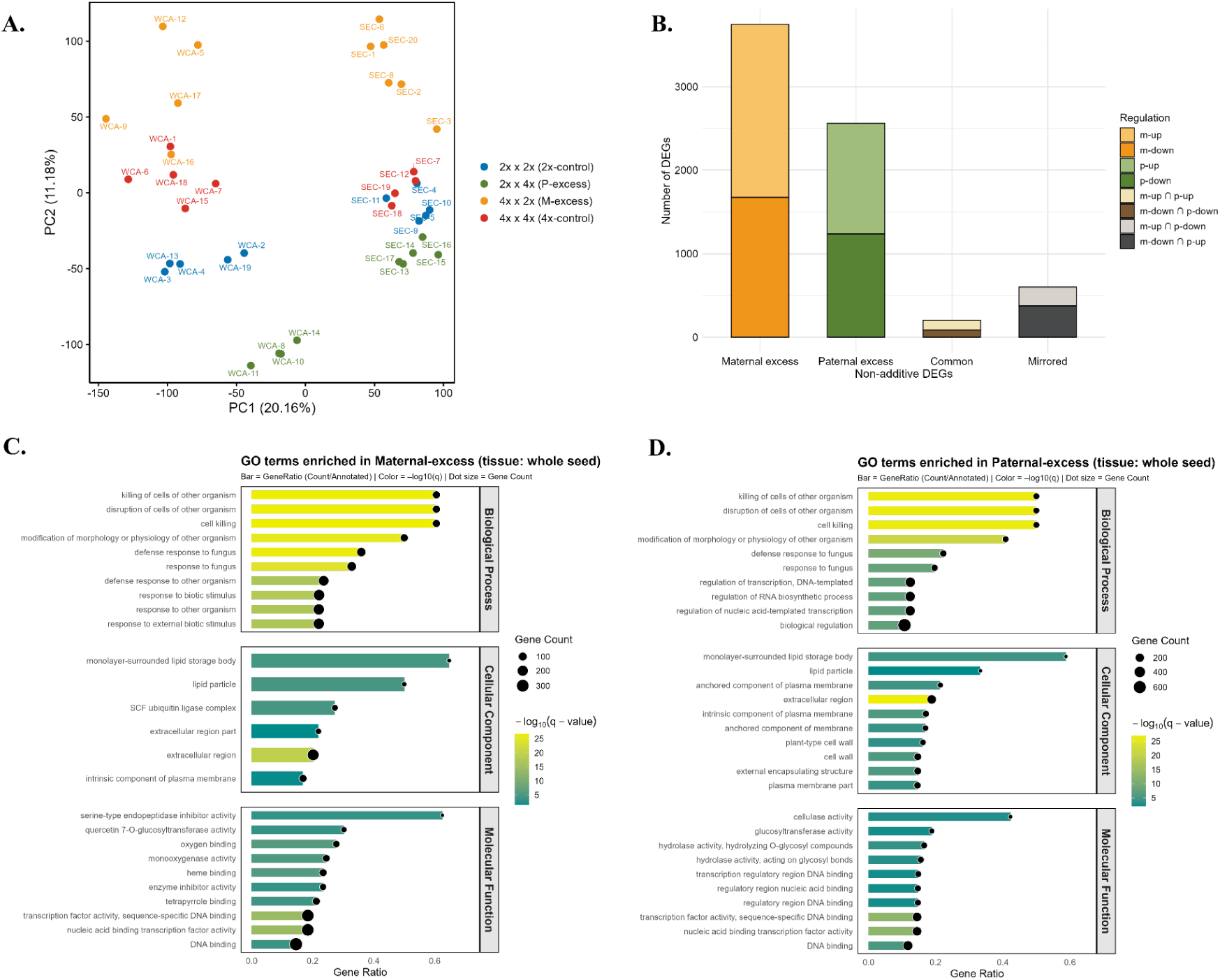
Overview of global transcriptomic analyses in interploidy hybrid seeds of *A. arenosa.* A. Principal component analysis (PCA) of whole-seed transcriptomes of 39 samples based on Transcripts per million (TPM) counts. B. Number of non-additively expressed genes in maternal- and paternal-excess crosses. C. Gene Ontology (GO) enrichment analysis of non-additive genes in maternal-excess crosses. Please note cellular component and molecular functions in maternal-excess were not significant enough. D. GO enrichment analysis of non-additive genes in paternal-excess crosses, across biological process, cellular component, and molecular function categories. In (C, D), dot size indicates gene count, color indicates q-value (FDR-adjusted p-value), and gene ratio denotes the proportion of input genes mapped to each term.

We then performed a differential expression analysis (DEA) comparing maternal- and paternal-excess hybrids with both diploid and tetraploid controls, focusing in particular on genes with non-additive expression in the hybrids. In order to leverage the full power of our dataset, we analysed all data (39 samples in total) together, accounting for the effect of the contact zone by considering it an additional factor in the DEA, yet focusing our interpretations on genes consistently differentially expressed in the focal contrast given by the crossing treatment. Maternal-excess (4x × 2x) hybrids showed a higher total number of non-additive differentially expressed genes, DEGs (n = 3744, 2071 up-regulated and 1673 down-regulated genes) compared to paternal-excess (2x × 4x; n = 2561 genes, 1324 up-regulated and 1237 down-regulated) (Fig. 1B, Supp Datasets 1-2).

To investigate the biological functions of these non-additive genes, we performed Gene Ontology (GO) enrichment analysis combining up- and down-regulated genes (Supp Datasets 3-4) because this reflects the biological reality of a transcriptional network with positive & negative expression relationships between genes. In maternal-excess hybrids (4x × 2x), GO enrichment of DEGs highlighted a strong overrepresentation of defense-related biological processes such as ‘response to fungus’, ‘defense response to other organism’, ‘disruption of cells of other organism’, and ‘cell killing’ (Fig. 1C). In paternal-excess hybrids (2x × 4x), GO enrichment (Fig. 1D) again revealed pathogen-related terms such as ‘response to fungus’ and ‘disruption of cells of other organism’. Additionally, cellular component terms such as ‘extracellular region’ and ‘plasma membrane part’, and molecular function terms like ‘hydrolase activity’ and ‘transcription factor activity’ were enriched.

### Disturbed expression of imprinted genes as a hallmark of interploidy hybrids

Imprinted genes have been causally linked to the triploid block in related species *A. thaliana* (Kradolfer et al., 2013; Wolff et al., 2015), and to test for their role in triploid block of *A. arenosa*, we first identified imprinted genes in diploid *A. arenosa* (47 MEGs, 49 PEGs; Supp Dataset 5). Orthologs of several of these genes are known to be imprinted in *A. thaliana*, such as *FIS2*, *HDG3* or *PEG5* (Supp Dataset 5; Luo et al., 2000; Jullien et al., 2006; Pignatta et al., 2018; Hornslien et al., 2019). To explore the expression patterns of imprinted genes in triploid seeds, we first performed a PCA using only imprinted gene TPM-normalized expression (Fig. 2A). Samples predominantly clustered by cross type this time rather than by contact zone, consistent with PERMANOVA results showing that cross type explained a larger proportion of variance (R² = 0.57) compared to contact zone (R² = 0.16), all statistically significant (p = 0.001) (Supp Table 1). In addition, we found a significant overrepresentation of both maternally (MEGs) and paternally (PEGs) expressed genes among the non-additive downregulated genes in triploid hybrid seeds (Fig. 2B). In particular, we observed a reciprocal pattern where PEGs were particularly downregulated in maternal-excess seeds (odds ratio > 10; Fig. 2B) and MEGs were particularly downregulated in paternal-excess seeds (odds ratio > 10).

**Fig. 2.**
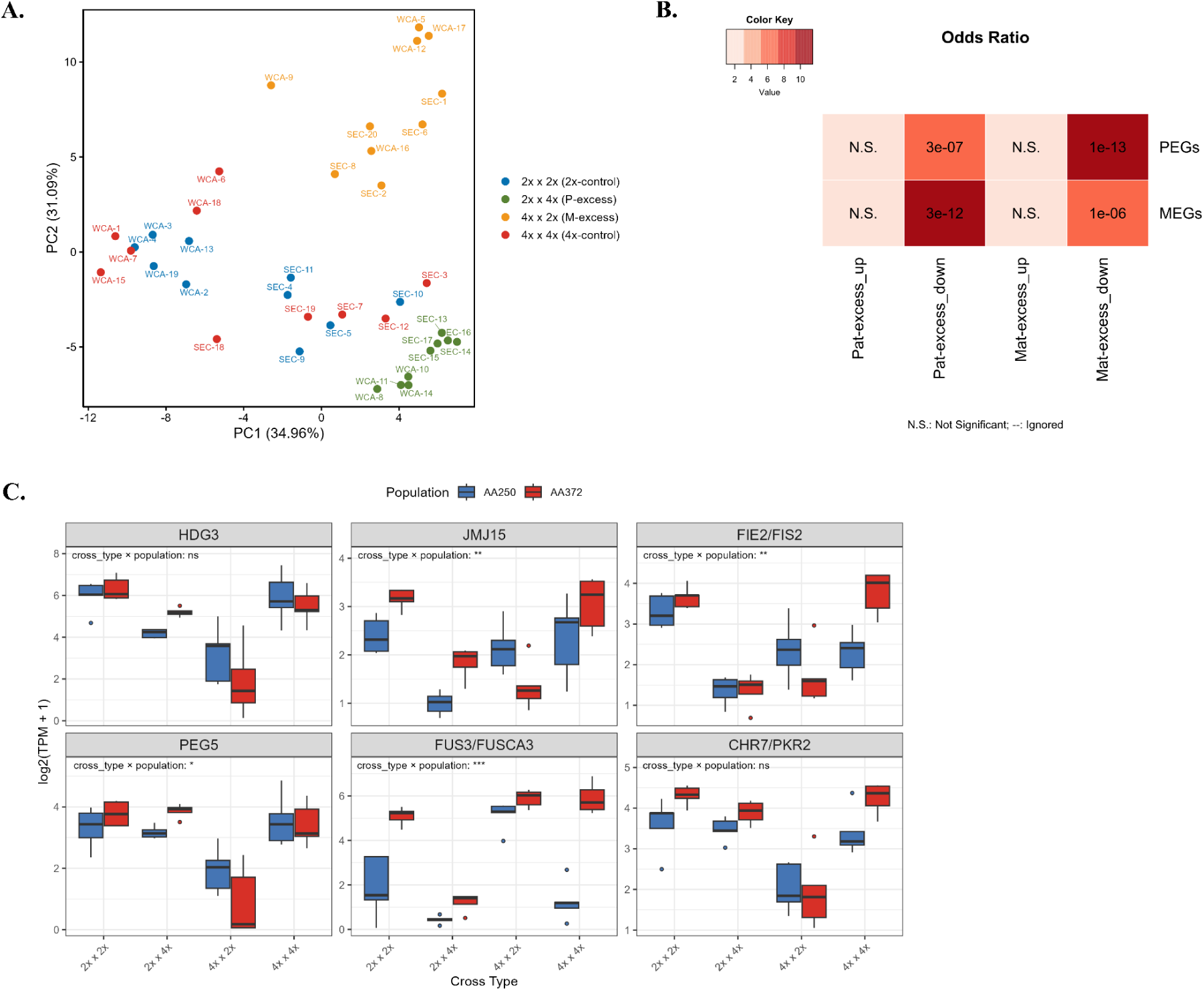
Expression analysis of imprinted genes in the intcrploidy hybrid seeds. **A.** PCA of transcript levels for the 96 imprinted genes inferred for diploid *A. arenosa* in this study across all samples. **B.** Odds ratio heatmap showing enrichment of maternally (MEGs) and paternally (l’EGs) expressed genes among up- and down-regulated genes in maternal- and paternal-excess crosses. Color scale indicates odds ratio; values represent p-values. N.S. = not significant. **C.** Expression patterns of a subset of imprinted genes across reciprocal interploidy crosses in two contact zones, SEC and WCA.

To further investigate the role of imprinted genes in *A. arenosa* triploid block, we examined the expression patterns of six imprinted genes (IGs) with a known or potential link to the triploid block in *A. thaliana* (Fig. 2C): *g15241* (*HDG3* – its imprinting affects endosperm cellularization; Pignatta et al., 2018), *g15720* (*PEG5* –misregulated in triploid seeds; Schatlowski et al., 2014), *g26034* (*PKR2* – mutating this gene partially suppresses the triploid block; Huang et al., 2017), *g15625* (*FIS2* – PRC2 component, its mutation phenocopies paternal excess; Chaudhury et al., 1997), *g15534* (*JMJ15* – histone demethylase; Hsieh et al., 2010) and *g17591* (*FUS3* – repressed by PRC2, its ectopic expression phenocopies paternal excess; Wu et al., 2020). In the maternal-excess (4x × 2x) cross, strong downregulation of several imprinted genes was observed relative to controls. This effect was especially pronounced for *HDG3*, *PEG5* and *PKR2*. These genes also showed relatively similar expression between diploid and tetraploid controls, suggesting a response specifically triggered by the maternal genomic excess. In contrast, *JMJ15* and *FIS2* showed down-regulation in both the maternal-excess and paternal-excess crosses. *FUS3* showed a mirrored expression pattern, being downregulated in paternal-excess and upregulated in maternal-excess. While some imprinted genes showed similar cross-type-specific expression across contact zones, cross type × contact zone interaction (F = 5.63, p = 0.00098) is significant, indicating lineage-specific differences in gene expression (Supp Table 2).

### Tissue-specific misregulation of gene expression in interploidy hybrids

To explore how different seed tissues contribute to transcriptional disruption in interploidy hybrids, we examined gene expression patterns in the embryo, endosperm, and seed coat separately, using a tissue deconvolution approach (see Methods section for details). We found significantly different relative tissue compositions between cross directions (Fig. S1), corresponding with phenotypic defects of hybrid seeds. In paternal-excess hybrid seeds (2x × 4x), the majority of non-additive gene expression was found in the endosperm and seed coat (N = 509 DEGs in each), and less so in the embryo (N = 190 DEGs) (Fig. 3A, Supp Datasets 6-8). In contrast, maternal-excess seeds (4x × 2x) showed a similar number of non-additively expressed genes in all tissues (n = 447, 471, and 663 for embryo, seed coat, and endosperm, respectively). In the maternal-excess cross—marked by a precocious endosperm cellularisation, embryo genes were predominantly upregulated, whereas endosperm and seed coat genes were largely downregulated. In the paternal-excess cross—characterized by delayed or arrested endosperm cellularization, seed coat genes showed mainly upregulation, while both embryo and endosperm genes were generally downregulated.

**Fig. 3.**
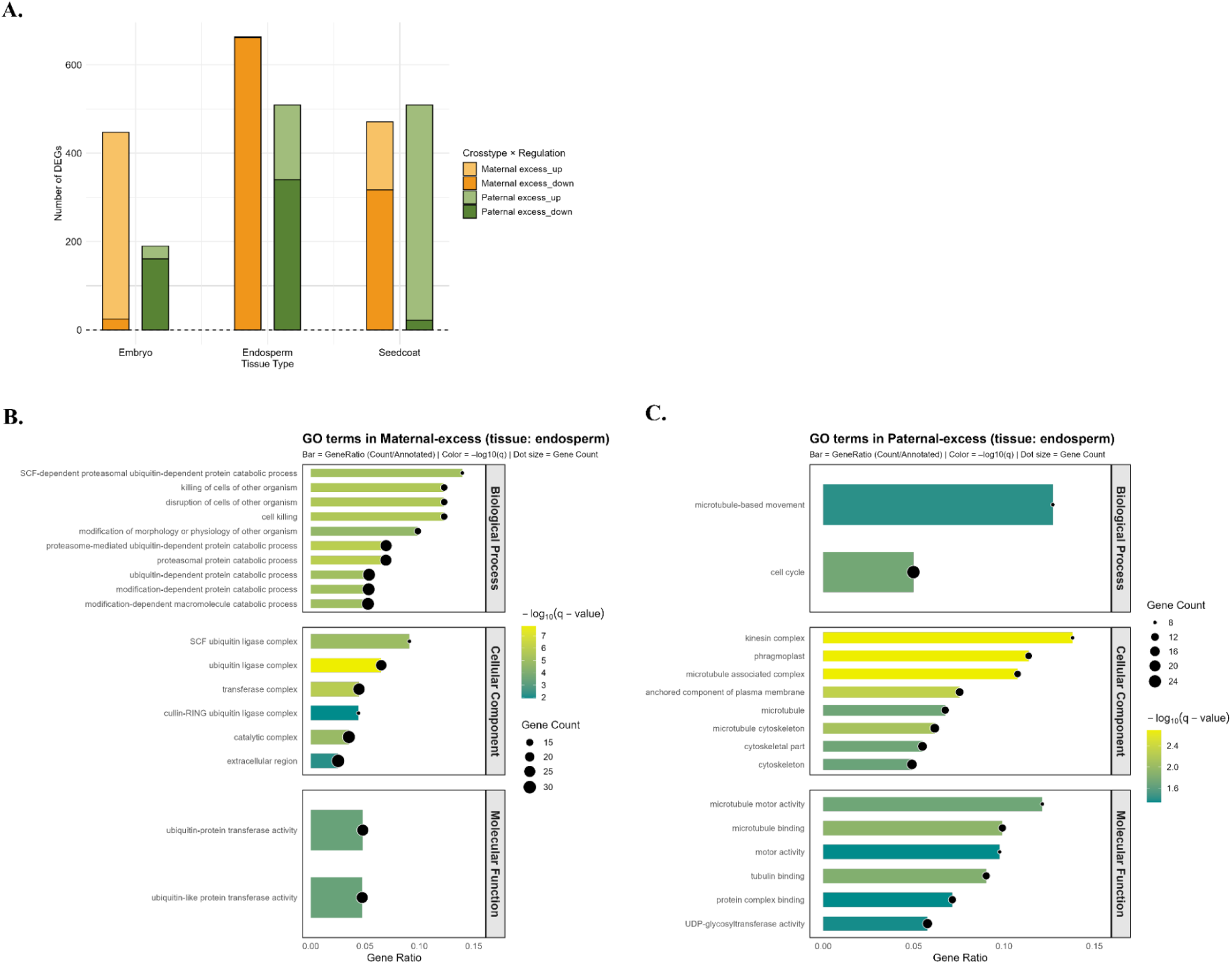
Tissue-specific responses to interploidy hybridization. **A.** Proportion of non-additively expressed genes across embryo, endosperm, and seed coat in maternal- and paternal-excess crosses. **(B-C).** GO enrichment of non-additive genes in endosperm for maternal- and paternal-excess crosses respectively. Dot size indicates gene count, color indicates q-value (FDR-adjusted p-value), and gene ratio denotes the proportion of input genes mapped to each term.

We then ran GO enrichment analyses separately for each tissue and cross type (Fig. 3B-C, Fig. 4A-D). In maternal-excess embryos, non-additive genes were enriched for lipid and transport-related terms, including ‘lipid particle’, ‘storage vacuole’, and ‘response to stress’ (Fig. 4A). Similarly, in paternal-excess embryos, enriched GO categories included ‘lipid particle’, ‘lipid storage’, and ‘nutrient reservoir activity’ (Fig. 4B), indicating that nutrient and metabolic pathways are commonly affected in embryo of both cross types.

**Fig. 4.**
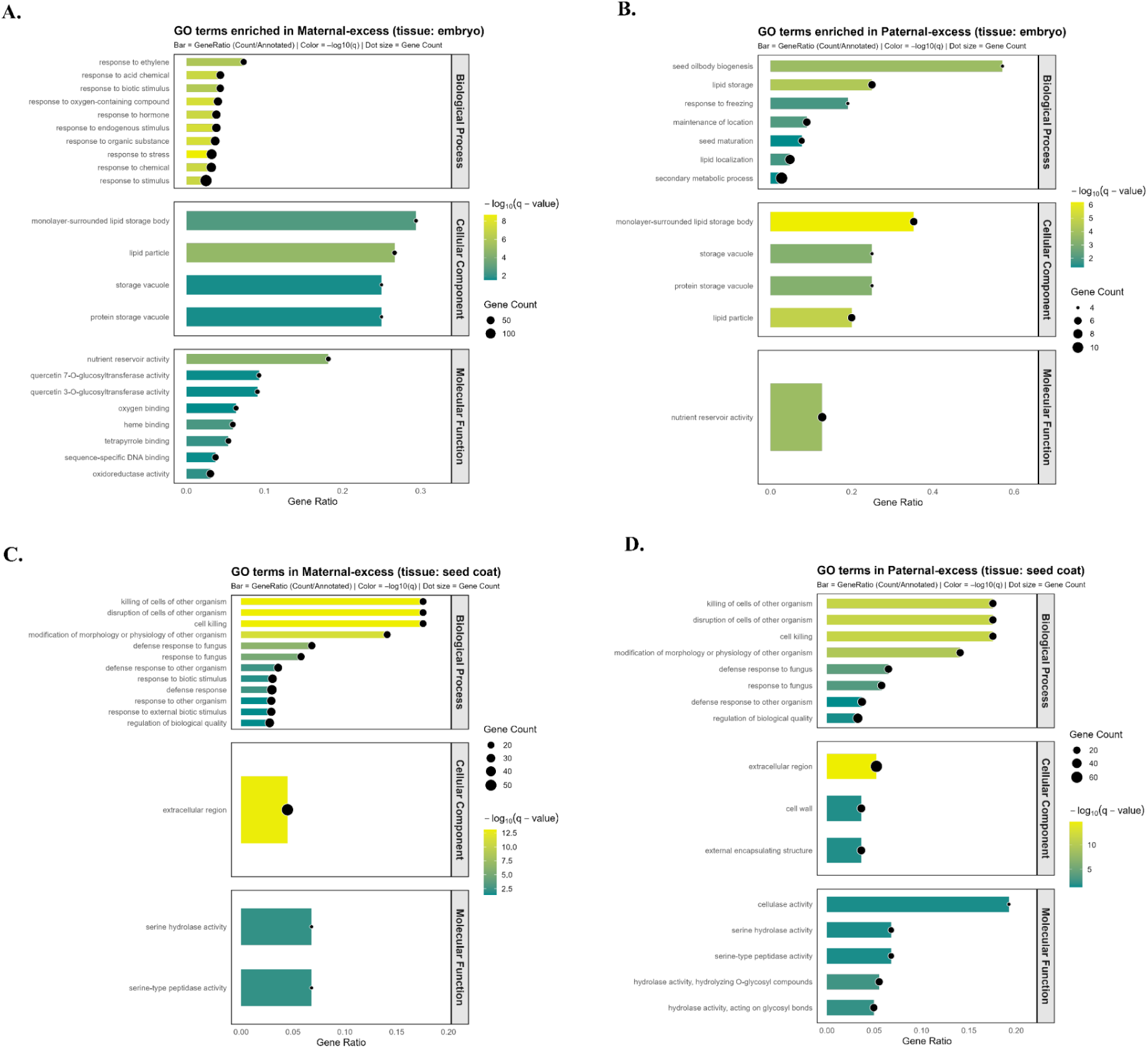
Tissue-specific responses to interploidy hybridization. **(A-D). GO** enrichment of non-additive genes in embryo **(A, B)** and seed coat **(C, D)** for maternal- and paternal-excess crosses respectively. Dot size indicares gene gene count, color indicates q-value (FDR-adjusted p-value), and gene ratio denotes the proportion of input genes mapped to each term.

In the endosperm, maternal-excess DEGs were significantly enriched for biotic stress and programmed cell death processes, including ‘killing of cells of other organism’ and ‘ubiquitin-dependent protein catabolic process’ (Fig. 3B). By contrast, paternal-excess DEGs were enriched for cell-cycle and structural processes such as ‘cell cycle’, ‘microtubule-based movement’, ‘cytoskeleton’, and ‘motor activity’ (Fig. 3C).

In the seed coat, non-additive genes from both maternal- and paternal-excess crosses were enriched for defense and stress-related terms, including ‘defense response’, ‘response to fungus’, ‘extracellular region’, and ‘serine hydrolase activity’ (Fig. 4C–D).

### Defense-like response is a conserved misregulated pathway in triploid seeds

Given the recurrence of enrichment in defense response GO terms, we wondered whether this pathway is also misregulated in triploid seeds of other species. For this purpose, we performed GO enrichment analyses on triploid seed DEGs identified in currently available triploid block studies: *A. thaliana* (paternal excess in Schatlowski et al., 2014: Ler x osd1, dissected endosperm; maternal excess in Tiwari et al., 2010: whole siliques), *Brassica rapa* (paternal excess in Stoute et al., 2012; whole seed) and *Oryza sativa* (maternal and paternal excess Wang et al., 2018; dissected endosperm). For paternal excess seeds, we found that 10 GO terms (Biological Processes) were commonly enriched in the paternal excess seeds of all species. This was more than expected by chance (SuperExactTest, P = 2.99e-08; Fig. 5A). Beyond expected terms like polysaccharide metabolism or cell wall biogenesis, the majority of GO terms were related to defense response (Fig. 5B). For maternal excess seeds, *B. rapa* data are not available, so we performed a three-way comparison, rice, *A. thaliana* and *A. arenosa*. A similar pattern was observed, i.e. 23 GO terms (BP) being enriched in all three species, among which defense response, more than expected by chance (SuperExactTest, P = 2.19e-13; Fig. S2).

**Fig. 5.**
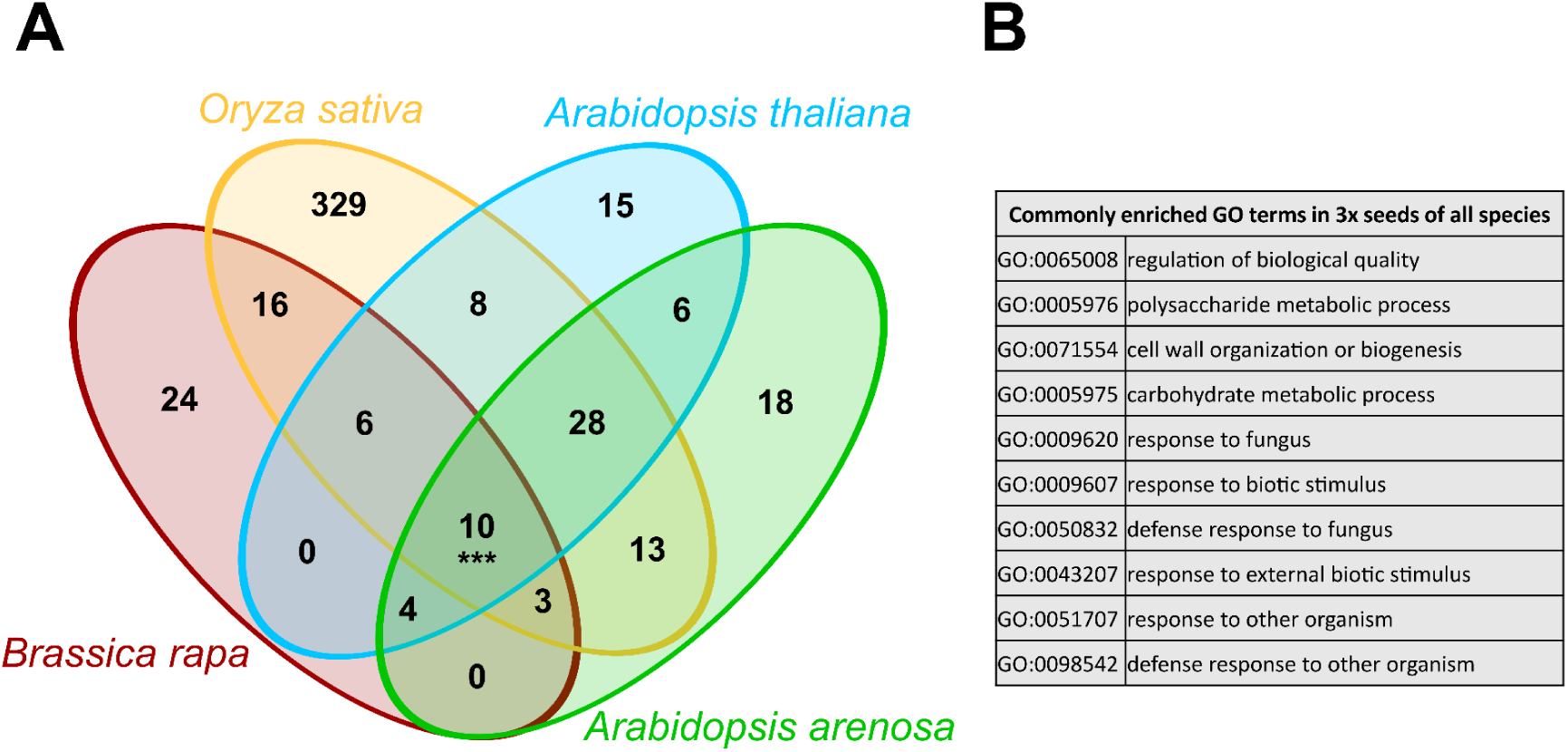
Functional pathways affected in paternal excess seeds across species studied to date *(Arabidopsis thaliana, Brassica rapa, Oryza sativa* and *A. arenosa* [this study]). **A.** Venn diagram showing the overlap of significantly enriched GO terms (Biological Process) in differentially expressed genes between paternal excess seeds and parents. *** indicates a significant overlap as tested with a SuperExact test (P = 2.99e-08). **B.** List of the 10 commonly enriched GO terms across all species and their description.

We further investigated the genes flagged with the defense-related GO terms to get a deeper insight into this phenomenon. Strikingly, every single gene with such annotation belonged to the defensin-like gene family, which are small cysteine-rich extracellular proteins, and only a few have been genetically characterized. These few genes were shown to be involved in cell-cell signalling related to sexual reproduction (Supp. DataSet S9).

## DISCUSSION

In this study, we investigated the transcriptomic response underlying triploid seed failure in *Arabidopsis arenosa*, a close relative of *A. thaliana* in which the triploid block acts in natural populations. We leveraged natural variation in order to have a representative picture of the species across its range, surveying genomic imprinting in 14 diploid populations, and monitoring the triploid block molecular response in two out the three known natural contact zones between diploids and tetraploids with distinct evolutionary history (Morgan et al., 2020, 2021). Within this evolutionarily relevant context, we identified known and also new molecular pathways involved in the triploid block. Our results, however, bear consequences beyond this species, bringing new insights on tissue-specific transcriptomic patterns and unraveling triploid block pathways conserved across all species studied so far.

### Imprinted genes, usual suspects

Despite a similar extent of misregulated genes, paternal-excess and maternal-excess seeds differ in their lethality rate and their phenotype, including the timing of endosperm cellularization and their shape and size (Morgan et al., 2021). This parent-of-origin phenomenon suggests genomic imprinting as an underlying molecular mechanism (Haig & Westoby, 1991), a mechanism that has been causally linked to the triploid block in *A. thaliana* (Kradolfer et al., 2013; Wolff et al., 2015). In this study, we found that the expression pattern of imprinted genes is a defining feature of maternal and paternal excess seeds, allowing us to discriminate these seeds from controls and from each other beyond population variation, which was otherwise the major source of expression variation across samples. Moreover, imprinted genes were preferentially misregulated in both maternal-excess and paternal-excess seeds. While this is not causal evidence, it is in line with previous studies showing that imprinted genes are misregulated in triploid seeds in *A. thaliana* and rice (Tiwari et al., 2010; Zhang et al., 2016). The cause of this misregulation is partially understood. Studies have shown that the imprinting status of these genes can be affected (Jullien & Berger, 2010; Zhang et al., 2016), which would then affect their general expression levels. In our study, we found that the maternally expressed *FIS2*, a master regulator of genomic imprinting (Batista & Köhler, 2020), was downregulated in both paternal and maternal-excess seeds. This is likely to have a snowball effect on both the expression level and the imprinting status of other imprinted genes. In addition, the small RNAs load in pollen has been causally linked to the lethality of paternal-excess seeds (Martinez et al., 2018; Borges et al., 2018), likely due to their epigenetic regulation of imprinted genes and the subsequent impact on expression levels and imprinting status of these genes. In this context, it would be interesting to investigate how small RNAs are involved in triploid seed defects in *A. arenosa*, in particular in the reduced viability of maternal-excess seeds that show natural phenotypic variation among populations (Morgan et al. 2021).

### Genes encoding small cysteine-rich proteins, overlooked yet conserved targets of triploid seed misregulation across species

We observed consistent enrichment of GO categories linked to stress and immunity, such as “response to stress”, “defense response to other organism”, and “cell killing”, among mis-regulated genes in interploidy hybrids. This pattern held for both paternal-excess (2x × 4x) and maternal-excess seeds. While this first suggested to us a potential pathogen infection during our experiments, this was easily ruled out by the fact that all types of crosses (including controls) were done on the same group of plants and were performed in parallel across the flowering period, avoiding any batch effect. Instead, by comparing our data with previous works in *A. thaliana*, *B. rapa* and *O. sativa* (Tiwari et al., 2010; Schatlowski et al., 2012; Stoute et al., 2012; Wang et al., 2018), we realized that these GO terms are systematically enriched in all studies and for both paternal and maternal excess seeds compared to parents. Surprisingly, none of these studies discuss this observation and its implication for the triploid block. Nevertheless, a study on *A. thaliana* × *A. arenosa* interspecific hybrid seeds found similarly over-represented immune response functions among misregulated genes (Burkart-Waco et al., 2013). The authors ruled out a potential auto-immune response as shown for other interspecific incompatibilities (Bomblies et al., 2007), and instead proposed that an immune response *per se* may not be happening. Instead, Burkart-Waco et al. proposed that genes annotated as “immune response” include cysteine-rich proteins, which are also important for cell-cell communication, notably between the different compartments of the seed (seed coat, endosperm and embryo; Marshall et al., 2011). This is supported by our data, showing that all genes tagged as defense-related, with no exception, belonged to the defensin-like/small cysteine-rich protein family. How these genes are connected to the triploid block remains unclear however. Most of them are functionally uncharacterized with no mutant described. For the ones that have been studied, they appear to be involved in cell-cell communication during reproduction, such as *SALVAGER1A* being produced in the central cell and attracting pollen tubes (Meng et al., 2023). We thus speculate that this triggered immune-like response is a result of the desynchronized development of the seed compartments – seed coat, embryo and endosperm – that happens during hybrid seed failure (Lafon-Placette & Köhler, 2014). The role of immune-like genes remains nevertheless unclear, which calls for further research to better understand their role in the triploid block.

### The characterization of tissue-specific responses provides a broader picture of the triploid block

Tissue-specific transcriptomic analyses revealed distinct yet interrelated patterns of transcriptional disruption across the endosperm, embryo, and seed coat in interploidy hybrid seeds. It is important to note that we used a deconvolution approach (Newman et al., 2019), inferring tissue specific expression from whole-seed transcriptomics in triploid seeds based on the transcriptome of actually isolated seed coat, embryo and endosperm for a subset of diploid samples. This does not replace tissue isolation and sequencing for all samples, but deconvolution has been shown to robustly reflect tissue specific expression patterns (Newman et al., 2019).

Disruption of endosperm cellularisation is a hallmark of the triploid block and a central feature of interploidy hybrid seed failure across diverse species, including *A. thaliana*, *B. napus*, and rice (Scott et al., 1998; Erilova et al., 2009; Rong et al., 2021; Wang et al., 2018; Tonosaki et al., 2020). This process depends on tightly regulated cytoskeletal organization and cell wall biosynthesis (Li et al., 2022). Consistently, we found that one of the only GO terms enriched in paternal excess seed DEGs across *A. arenosa*, *A. thaliana, B. napus* and rice involved cell wall biogenesis. Our study showed misregulation of genes involved in cytoskeleton organization and microtubule-based movement specifically in the endosperm of paternal-excess *A. arenosa* seeds. These are important components of mitosis, but also of cell wall formation (Yadav, 2025), likely contributing to the endosperm cellularization defects underlying seed failure in paternal-excess seeds. In contrast, the transcriptome of maternal-excess hybrid endosperm exhibited enriched functions in defense response, as well as proteasomal degradation and ubiquitination of proteins. The latter pathway has been associated with the regulation of seed size (Li & Li, 2014). Moreover, mutations of specific components of ubiquitin-mediated protein degradation, *CUL3A* and *CUL3B*, led to delayed or failed endosperm cellularization and embryo arrest, suggesting an important yet not fully clear role of this pathway in normal endosperm and embryo development (Thomann et al., 2005). How this pathway impacts maternal-excess seeds development remains even less clear, requiring further research.

In both maternal- and paternal-excess hybrids, misregulated genes in the embryo displayed significant enrichment for GO categories linked to lipid metabolism and nutrient storage, including ‘lipid particle’, ‘lipid storage’, ‘storage vacuole’, and ‘nutrient reservoir activity’. In seeds, these categories are not simply markers of altered metabolism but are direct targets of the ABA-driven maturation network that coordinate reserve accumulation, desiccation tolerance, and dormancy during the late maturation phase (Baud et al., 2008; Santos-Mendoza et al., 2008). If the endosperm cannot supply enough nutrients or hormones, the embryo prematurely initiates maturation-like programs, likely as a compensatory response to compromised nutrient provisioning or hormonal cues from the dysfunctional endosperm. A similar shift has been observed in *A. thaliana* triploid embryos, where gene expression shifts toward lipid metabolism and desiccation tolerance resemble late seed maturation (Xu et al., 2023). These embryo-level responses likely reflect an evolutionarily conserved survival mechanism triggered by upstream failure in the endosperm, positioning the embryo as a secondary but critical site of transcriptional disruption during hybrid seed breakdown.

The seed coat showed robust enrichment for defense-related and extracellular signaling GO categories, such as ‘defense response’, ‘response to fungus’, ‘serine hydrolase activity’, and ‘extracellular region’, in both maternal- and paternal-excess crosses, with particularly strong upregulation observed in paternal-excess hybrids. The endosperm DEGs also showed enrichment in these functions, unlike the embryo, ruling out a generalized auto-immune response in triploid seeds. Instead, as discussed in the previous section, this immune-like response may suggest a disturbed communication between the maternal seed coat and the endosperm, as proteins involved in cell-cell communication also play a role in the immune response (Marshall et al., 2011). Alternatively, the seed coat contains a mechano-sensing cell layer able to detect the mechanical pressure exerted by the growing endosperm, which is important for seed coat development (Creff et al., 2015). Mechanical pressure on cell walls exerted by growing pathogens is also a cue for defense response activation (Léger et al., 2022). An under- or overgrowing endosperm in maternal- and paternal-excess seeds respectively, and the subsequent abnormal mechanical pressure on the seedcoat could thus affect immune responses. Importantly, mechanical pressure may also happen the other way around, from seed coat to endosperm, and regulate endosperm development. Recent work has highlighted the maternal regulatory role of the seed coat in *A. thaliana* via transcription factors like *TRANSPARENT TESTA 8* (TT8), whose mutation allows the rescue of paternal-excess seeds putatively by reducing seed coat expansion and thus buffering endosperm overgrowth (Zumajo-Cardona et al., 2023). Our data suggest that the seed coat senses endosperm imbalanced growth, potentially through mechano-sensing triggering immune-like responses. The repeated enrichment of ‘extracellular region’ and defense-related genes suggests that disrupted communication between the seed coat and endosperm contributes to hybrid seed failure in interploidy crosses.

These findings reinforce the importance of inter-tissue communication in seed development and highlight its breakdown as a central mechanism of postzygotic reproductive isolation in interploidy hybrids.

## ACKNOWLEDGEMENTS

The authors acknowledge the special support of V. Vlčková for her assistance with the laboratory cultivations and crossing experiments. Flow cytometry and growth chamber facilities were provided by the Department of Botany, Charles University in Prague, Czech Republic. We also acknowledge Duarte D. Figueiredo and Ana Marcela Florez-Reuda for their insightful discussions during the preliminary stages of data analysis.

## COMPETING INTERESTS

The authors declare that they have no conflict of interest.

## AUTHOR CONTRIBUTIONS

S.S. conducted the crossing experiment, sample preparation and RNA extraction, data analysis, and interpretation related to the triploid block experiment. V.C. and A.Pr. prepared samples and performed experiments for imprinted gene study. M.K. performed the identification of imprinted genes and genome annotation. F.K., C.L.P. and A.Pe. contributed to the experimental design and data interpretation. F.K. and C.L.P. also conceived, designed, and supervised the overall project. S.S., F.K. and C.L.P. jointly wrote the manuscript. All authors reviewed and approved the final version for publication.

## DATA AVAILABILITY

RNA and DNA data for the imprinting analysis are available in ENA with the project accession number PRJEB79247. RNA-Seq data generated for the triploid block experiment have been deposited in the NCBI Sequence Read Archive (SRA) under BioProject accession number PRJNA1433134.

All supplementary datasets, figures and tables are provided in the Supporting Information of this article.

## Supporting Information

**Supplementary Figure 1.** Tissue-specific deconvolution.

**Supplementary Figure 2.** Functional pathways affected in maternal excess seeds across species studied to date.

**Supplementary Figure 3.** Representation of population locations and crossing design.

**Supplementary Table 1.** Results of the PERMANOVA test, based either on normalized RNA-Seq readcounts of all genes or of imprinted genes.

**Supplementary Table 2.** Summary statistics of imprinted gene expression from GLM analysis.

**Supplementary Table 3.** Details of the populations studied in this article.

**Supplementary Table 4.** Crossing design for the imprinted genes survey.

**Supplementary Table 5.** Crossing design for the triploid block study.

**Supplementary Dataset 1.** List of differentially expressed genes in the maternal excess cross.

**Supplementary Dataset 2.** List of differentially expressed genes in the paternal excess cross.

**Supplementary Dataset 3.** Enriched GO terms among DEGs in the maternal excess cross.

**Supplementary Dataset 4.** Enriched GO terms among DEGs in the paternal excess cross.

**Supplementary Dataset 5.** List of imprinted genes identified in *A. arenosa* and their annotation in *A. thaliana*.

**Supplementary Dataset 6.** List of differentially expressed genes from the embryo based on deconvolution.

**Supplementary Dataset 7.** List of differentially expressed genes from the endosperm based on deconvolution.

**Supplementary Dataset 8.** List of differentially expressed genes from the seed coat based on deconvolution.

**Supplementary Dataset 9.** List of DEGs (paternal excess) flagged with the GO term “cell killing”.

**Supplementary Dataset 10.** List of *A. arenosa* genes and their identified *A. thaliana* orthologues.

## Notes

### Competing Interest Statement

The authors have declared no competing interest.

### Funding

- Czech Science Foundation

