## Supplementary figures&tables for "Transcriptomic insights into triploid seed failure in *Arabidopsis arenosa* natural populations"

A.

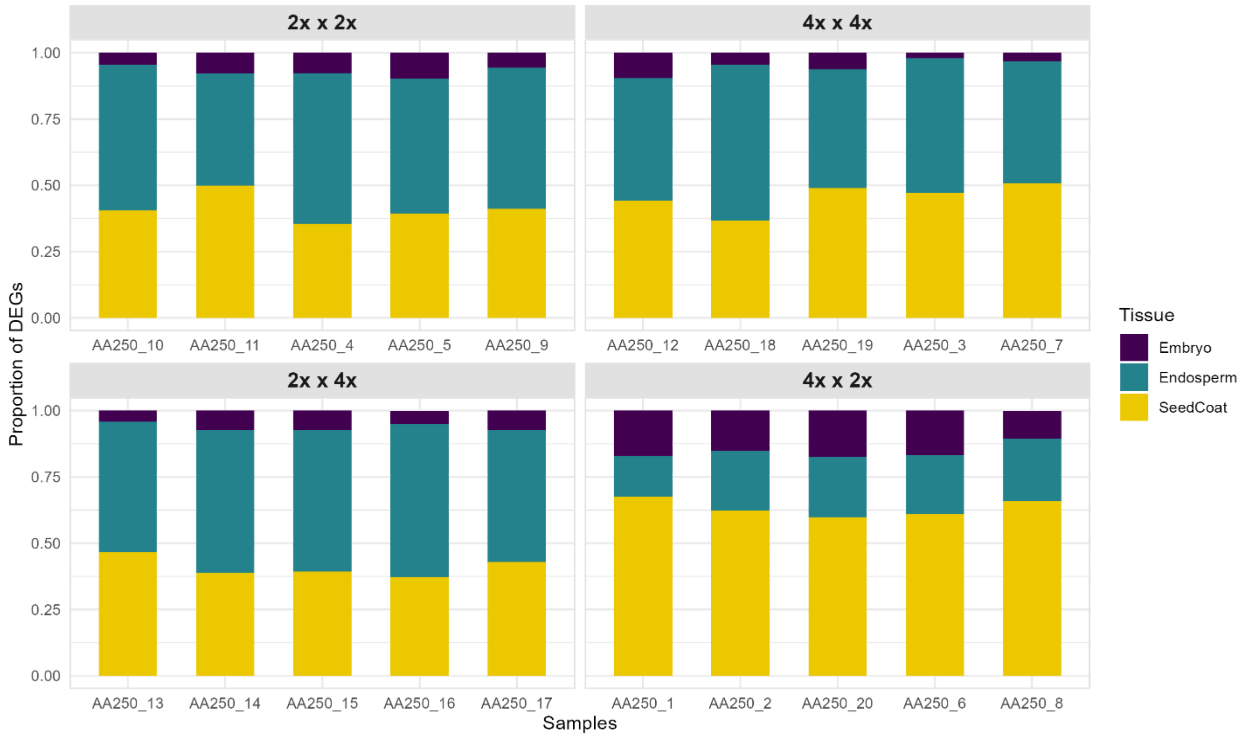


B.

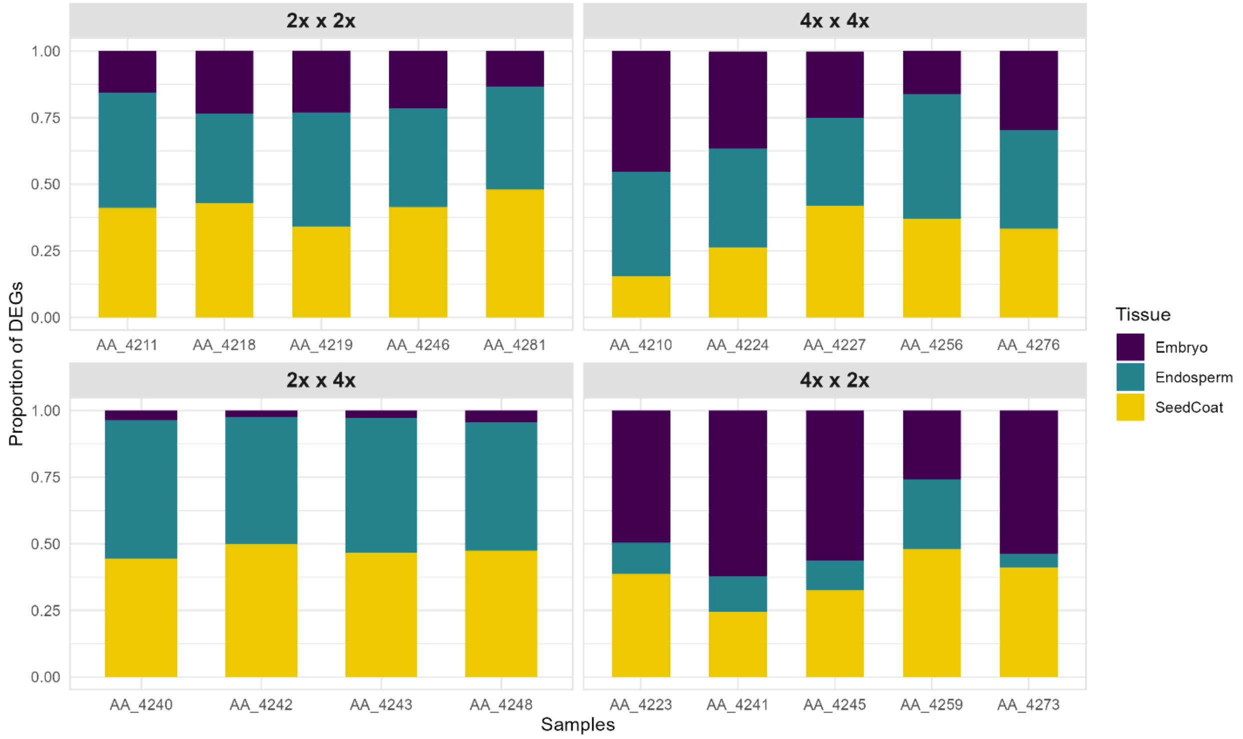


**Fig. S1** Tissue-specific deconvolution. **(A)** Estimated tissue proportions in SEC samples based on tissue deconvolution. **(B)** Estimated tissue proportions in WCA samples based on tissue deconvolution.

**A**

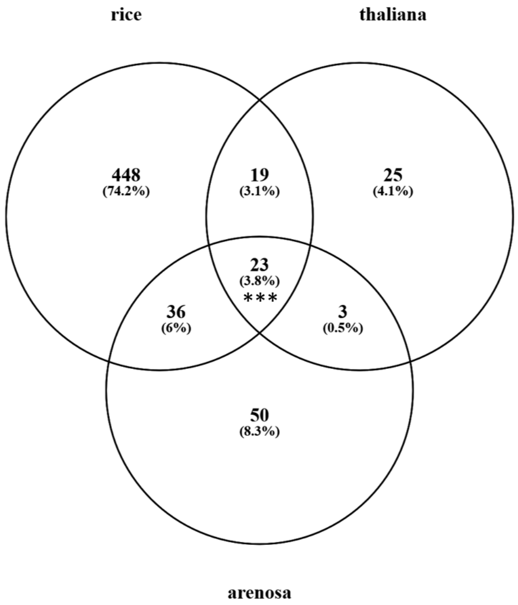


**B**

| Commonly enriched GO terms in 3x seeds of all species | |
|---|---|
| GO:0009607 | response to biotic stimulus |
| GO:0043207 | response to external biotic stimulus |
| GO:0051707 | response to other organism |
| GO:0098542 | defense response to other organism |
| GO:0006952 | defense response |
| GO:0009605 | response to external stimulus |
| GO:0051704 | multi-organism process |
| GO:0009719 | response to endogenous stimulus |
| GO:0010033 | response to organic substance |
| GO:0009725 | response to hormone |
| GO:0006950 | response to stress |
| GO:1901700 | response to oxygen-containing compound |
| GO:0001101 | response to acid chemical |
| GO:0042221 | response to chemical |
| GO:0050896 | response to stimulus |
| GO:0071495 | cellular response to endogenous stimulus |
| GO:0007568 | aging |
| GO:0032870 | cellular response to hormone stimulus |
| GO:0010150 | leaf senescence |
| GO:0010260 | organ senescence |
| GO:0009755 | hormone-mediated signaling pathway |
| GO:0070887 | cellular response to chemical stimulus |
| GO:0071310 | cellular response to organic substance |

**Fig. S2** Functional pathways affected in maternal excess seeds across species studied to date (*Arabidopsis thaliana*, *Oryza sativa,* and *A. arenosa* [this study]). **(A)** Venn diagram showing the overlap of significantly enriched GO terms (Biological Process) in differentially expressed genes between maternal-excess seeds and parents. *** indicates a significant overlap as tested with a SuperExact test (P = 2.19e-13). **(B)** List of the 23 commonly enriched GO terms across all species and their description.

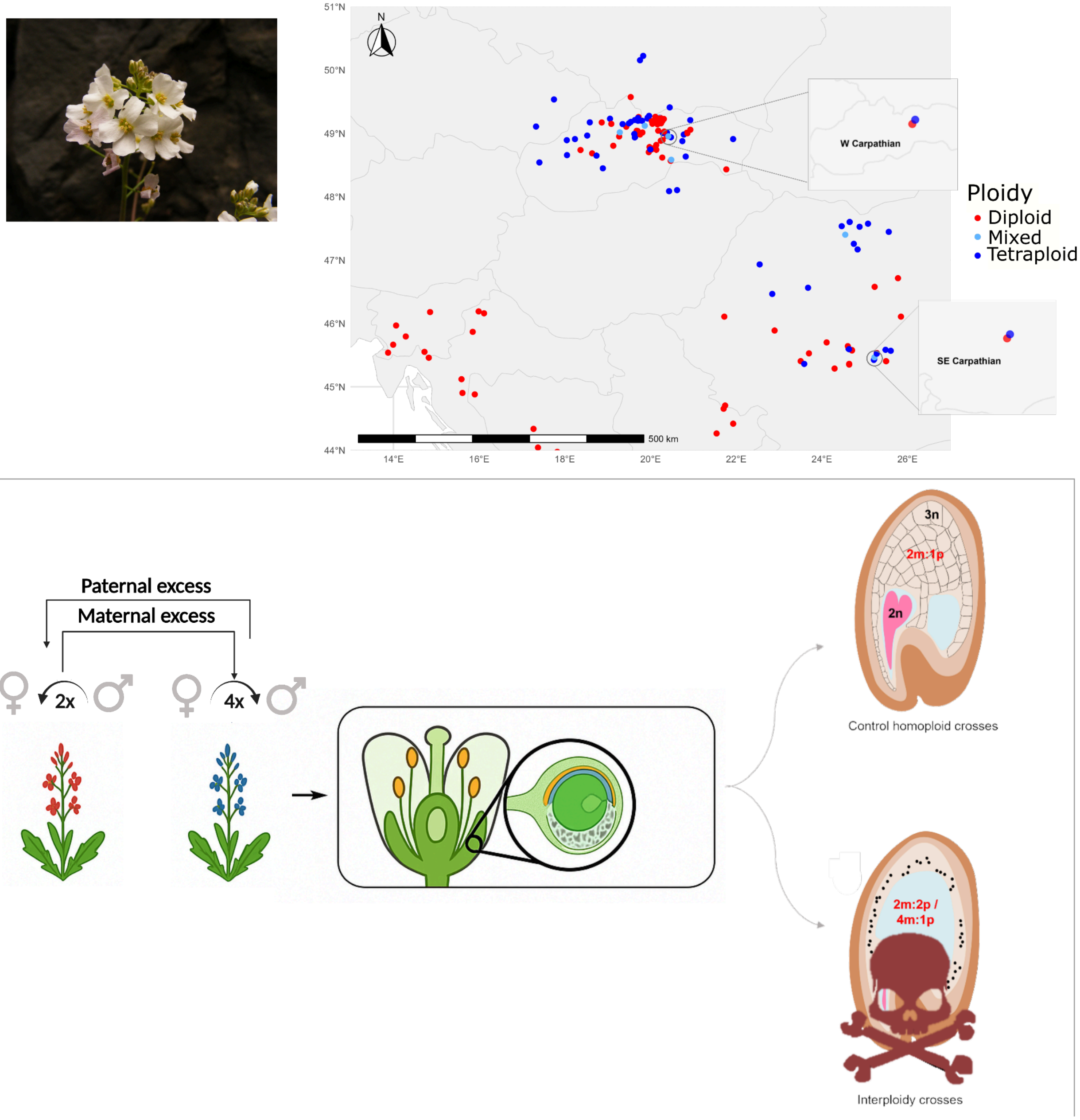


**Fig. S3** Map of *A. arenosa* diploid and tetraploid populations (top) and schematic representation of the crossing design (bottom).

**Table S1.** Results of the PERMANOVA test, based either on normalized RNA-Seq readcounts of all genes or of imprinted genes.

| Model | Df | SumOfSqs | R2 | F | Pr(>F) |
|---|---|---|---|---|---|
| *All genes* | | | | | |
| cross_type | 3 | 173928 | 0.29267 | 4.8272 | 0.001 |
| population | 1 | 148446 | 0.24979 | 12.319 | 0.001 |
| cross_type * population | 7 | 377691 | 0.63554 | 7.7223 | 0.001 |
| *Imprinted genes* | | | | | |
| cross_type | 3 | 2475.1 | 0.56996 | 15.462 | 0.001 |
| population | 1 | 713.7 | 0.16436 | 7.2773 | 0.001 |
| cross_type * population | 7 | 3541.8 | 0.81558 | 19.585 | 0.001 |

**Table S2.** Summary statistics of imprinted gene expression from GLM analysis.

| Term | F-value | p-value |
|---|---|---|
| cross_type | 20.55 | *** 8.79e-12 |
| population | 12.08 | ** 0.00061 |
| gene_id | 20.61 | *** <2e-16 |
| cross_type:population | 5.63 | ** 0.00098 |

**Table S3.** Details of the populations studied in this article.

| Population code | Ploidy | Lineages | Country | Latitude | Orientation | Longitude | Orientation | Experiment |
|---|---|---|---|---|---|---|---|---|
| AA025 | 2x | WCA | Slovakia | 49.17408333 | N | 18.8617 | E | Imprinted genes survey |
| AA034 | 2x | PA | Hungary | 47.458 | N | 18.924778 | E | Imprinted genes survey |
| AA038 | 2x | PA | Hungary | 46.09916667 | N | 18.233917 | E | Imprinted genes survey |
| AA054 | 2x | DI | Croatia | 44.90433333 | N | 15.6111 | E | Imprinted genes survey |
| AA084 | 2x | WCA | Slovakia | 49.162 | N | 20.154194 | E | Imprinted genes survey |
| AA090 | 2x | WCA | Slovakia | 49.20652778 | N | 20.215056 | E | Imprinted genes survey |
| AA106 | 2x | DI | Croatia | 46.16166667 | N | 16.115 | E | Imprinted genes survey |
| AA130 | 2x | DI | Bosnia and Herzegovina | 44.88180833 | N | 15.898817 | E | Imprinted genes survey |
| AA161 | 2x | DI | Slovenia | 45.795847 | N | 14.287934 | E | Imprinted genes survey |
| AA208 | 2x | WCA | Slovakia | 49.043514 | N | 20.180772 | E | Imprinted genes survey |
| AA250 | 2x | SEC | Romania | 45.53349 | N | 25.29145 | E | Triploid block |
| AA250 | 4x | SEC | Romania | 45.52998 | N | 25.26694 | E | Triploid block |
| AA347 | 2x | PA | Slovakia | 48.26694 | N | 19 | E | Imprinted genes survey |
| AA349 | 2x | PA | Hungary | 46.80667 | N | 17.43444 | E | Imprinted genes survey |
| AA372 | 2x | WCA | Slovakia | 48.96221 | N | 20.40265 | E | Imprinted genes survey + Triploid block |
| AA372 | 4x | WCA | Slovakia | 48.95614 | N | 20.41459 | E | Triploid block |
| AA433 | 2x | WCA | Slovakia | 48.82446 | N | 19.022602 | E | Imprinted genes survey |

**Table S4.** Crossing design for the imprinted genes survey.

| Imprintome | Maternal plant | | Paternal plant | |
|---|---|---|---|---|
| | Population code | Individual | Population code | Individual |
| 1 | AA208 | A | AA084 | A |
| | AA084 | A | AA208 | A |
| 2 | AA208 | B | AA084 | B |
| | AA084 | B | AA208 | B |
| 3 | AA372 | A | AA090 | A |
| | AA090 | A | AA372 | A |
| 4 | AA106 | A | AA130 | A |
| | AA130 | A | AA106 | A |
| 5 | AA034 | A | AA347 | A |
| | AA347 | A | AA034 | A |
| 6 | AA034 | B | AA347 | B |
| | AA347 | B | AA034 | B |
| 7 | AA084 | C | AA208 | C |
| | AA208 | C | AA084 | C |
| 8 | AA084 | D | AA208 | D |
| | AA208 | D | AA084 | D |
| 9 | AA054 | A | AA161 | A |
| | AA161 | A | AA054 | A |
| 10 | AA433 | A | AA025 | A |
| | AA025 | A | AA433 | A |
| 11 | AA038 | A | AA349 | A |
| | AA349 | A | AA038 | A |
| 12 | AA038 | B | AA349 | B |
| | AA349 | B | AA038 | B |
| 13 | AA038 | C | AA349 | C |
| | AA349 | C | AA038 | C |

**Table S5.** Crossing design for the triploid block study.

| Maternal plant | | | Paternal plant | | | |
|---|---|---|---|---|---|---|
| **Population** | **Ploidy** | **Individual** | **Population** | **Ploidy** | **Individual** | **Cross type** |
| AA250 | 2x | BC2 | AA250 | 2x | DQ3 | diploid control |
| AA250 | 2x | DZ1 | AA250 | 2x | DQ2 | diploid control |
| AA250 | 2x | DQ1 | AA250 | 2x | BC4 | diploid control |
| AA250 | 2x | DZ2 | AA250 | 2x | BC3 | diploid control |
| AA250 | 2x | BC1 | AA250 | 2x | DZ3 | diploid control |
| AA250 | 4x | BF1 | AA250 | 4x | CK2 | tetraploid control |
| AA250 | 4x | BF2 | AA250 | 4x | CC3 | tetraploid control |
| AA250 | 4x | CC4 | AA250 | 4x | CK3 | tetraploid control |
| AA250 | 4x | CK1 | AA250 | 4x | BF3 | tetraploid control |
| AA250 | 4x | CC1 | AA250 | 4x | BF4 | tetraploid control |
| AA250 | 2x | DZ2 | AA250 | 4x | CC3 | paternal excess |
| AA250 | 2x | BC1 | AA250 | 4x | BF4 | paternal excess |
| AA250 | 2x | DQ8 | AA250 | 4x | BF3 | paternal excess |
| AA250 | 2x | BC2 | AA250 | 4x | CK3 | paternal excess |
| AA250 | 2x | DZ1 | AA250 | 4x | CK2 | paternal excess |
| AA250 | 4x | CK6 | AA250 | 2x | BC4 | maternal excess |
| AA250 | 4x | BF2 | AA250 | 2x | BC3 | maternal excess |
| AA250 | 4x | CC4 | AA250 | 2x | DQ3 | maternal excess |
| AA250 | 4x | BF1 | AA250 | 2x | DQ2 | maternal excess |
| AA250 | 4x | CC1 | AA250 | 2x | DZ3 | maternal excess |
| AA372 | 2x | 258C | AA372 | 2x | 175C | diploid control |
| AA372 | 2x | 258B | AA372 | 2x | 175B | diploid control |
| AA372 | 2x | 258A | AA372 | 2x | 175A | diploid control |
| AA372 | 2x | 175E | AA372 | 2x | 258E | diploid control |
| AA372 | 2x | 175D | AA372 | 2x | 258D | diploid control |
| AA372 | 4x | 208C | AA372 | 4x | 206E | tetraploid control |
| AA372 | 4x | 206A | AA372 | 4x | 208A | tetraploid control |
| AA372 | 4x | 206B | AA372 | 4x | 208B | tetraploid control |
| AA372 | 4x | 206C | AA372 | 4x | 208A | tetraploid control |
| AA372 | 4x | 208D | AA372 | 4x | 206D | tetraploid control |
| AA372 | 2x | 258C | AA372 | 4x | 208A | paternal excess |
| AA372 | 2x | 175D | AA372 | 4x | 206D | paternal excess |
| AA372 | 2x | 175E | AA372 | 4x | 206E | paternal excess |
| AA372 | 2x | 258B | AA372 | 4x | 208B | paternal excess |
| AA372 | 2x | 258A | AA372 | 4x | 208A | paternal excess |
| AA372 | 4x | 206A | AA372 | 2x | 175A | maternal excess |
| AA372 | 4x | 206B | AA372 | 2x | 175B | maternal excess |
| AA372 | 4x | 206C | AA372 | 2x | 175C | maternal excess |
| AA372 | 4x | 208D | AA372 | 2x | 258D | maternal excess |
| AA372 | 4x | 208C | AA372 | 2x | 258E | maternal excess |
